# MAP Kinase inhibition reshapes tumor microenvironment of mouse pancreatic cancer by depleting anti-inflammatory macrophages

**DOI:** 10.1101/759779

**Authors:** Pietro Delfino, Christian Neander, Dea Filippini, Sabrina L. D’Agosto, Caterina Vicentini, Elena Fiorini, Francesca Lupo, Claudia Fiorini, Borislav Rusev, Gael D. Temgue Tane, Francesco De Sanctis, Michele Simbolo, Stefano Ugel, Vincenzo Bronte, Matteo Fassan, Rita T. Lawlor, Federico Boschi, Stefano Barbi, Aldo Scarpa, Phyllis F. Cheung, Jens T. Siveke, Vincenzo Corbo

## Abstract

The RAF/MEK/ERK (MAP Kinase) pathway is the index oncogenic signaling towards which many compounds have been developed and tested for the treatment of KRAS-driven cancers, including pancreatic ductal adenocarcinoma (PDA). Here, we explored the immunological changes induced by targeted MEK1/2 inhibition (MEKi) using trametinib in preclinical mouse models of PDA. We evaluated the dynamic changes in the immune contexture of mouse PDA upon MEKi using a multidimensional approach (mRNA analyses, flow cytometry, and immunophenotyping). Effect of MEKi on the viability and metabolism of macrophages was investigated *in vitro*. We showed that transcriptional signatures of MAP Kinase activation are enriched in aggressive human PDA subtype (squamous/basal-like/quasimesenchymal), while short term MEKi treatment in mouse PDA induced subtype switching. Integrative mRNA expression and immunophenotypic analyses showed that MEKi reshapes the immune landscape of PDA by depleting rather than reprogramming macrophages, while augmenting infiltration by neutrophils. Depletion of macrophages is observed early in the course of *in vivo* treatment and is at least partially due to their higher sensitivity to MEKi. Tumor-associated macrophages were consistently reported to interfere with gemcitabine uptake by PDA cells. Here, our *in vivo* studies show a superior antitumor activity upon combination of MEKi and gemcitabine using a sequential rather than simultaneous dosing protocol. Our results show that MEK inhibition induces a dramatic remodeling of the tumor microenvironment of mouse PDA through depletion of macrophages, which substantially improves the antitumor activity of gemcitabine.

## INTRODUCTION

Pancreatic ductal adenocarcinoma (PDA) has the lowest survival rate of all cancers (1). Current therapeutic strategies are mostly based on combination of chemotherapeutic agents, which provides intermittent response and modest survival benefit in the advanced disease setting (2,3). Results from recent clinical trials showed that immune-checkpoint based therapy is rarely active in PDA (4). The complex and heterogenous PDA microenvironment is reported to play a major role in mediating resistance to both chemotherapy and immune therapy (5). In human PDA stromal non-malignant cells often outnumber neoplastic cells, with fibroblasts and macrophages being the dominants cellular components (6).

Activating mutations of the GTPase *KRAS* are a nearly universal feature of PDA (7). Activated KRAS engages a multitude of pathways, including the RAF/MEK/ERK (MAP Kinase) pathway, which regulates cellular processes such as proliferation and cell survival (8). Preclinical mouse models show that the oncogenic KRAS-induced cytokine milieu likely contributes to the highly immunosuppressive microenvironment observed already at the preneoplastic PanIN stage (9). In addition to non-cell autonomous effects, KRAS activation and increased activity through the MAP Kinase pathway induce cell-intrinsic upregulation of PD-L1, thus inhibiting anti-tumor immunity (10). Several molecular classification systems have been proposed for PDA (7,11,12). Of the four subtypes of PDA described by Bailey et al. (7), two (immunogenic and squamous) revealed high expression of inflammatory-related transcripts, including a macrophage transcriptional signature that indicated worse patients’ survival (7). Furthermore, the squamous subtype is dominated by transcriptional signatures indicative of a highly immune suppressed microenvironment, and accordingly cytotoxic T cell signatures are absent (7).

When the neoplastic epithelium is experimentally or computationally microdissected away from surrounding stroma (11,12), two major PDA subtypes emerge from transcriptomic analysis, which are fairly aligned across the three proposed classifications (7,11,12): the squamous/quasimesenchymal/basal-like subtype and the pancreatic progenitor/classical subtype.

Depletion of myeloid cells/macrophages through genetic and pharmacological approaches have demonstrated promising for increasing efficacy of immune checkpoint inhibitors (13,14). Of these myeloid cell populations, tumor-associated macrophages play a relevant role in tumor progression (14), resistance to chemotherapy (15,16), and immune tolerance in PDA (17).

Despite decades of extensive efforts, KRAS has remained an undruggable target (18). Few drugs exist that target allele-specific mutations (e.g., KRAS p.G12C), which are poorly represented in PDA accounting for 1% of mutated *KRAS* (8). Therefore, most efforts have been focusing on the development of small molecules that hit downstream nodes of activated KRAS including the Ser/Thr kinases MEK1/2 (18). While several studies have evaluated efficacy of targeted therapies and identified cell-intrinsic mechanisms of resistance, a systematic characterization of the changes in the immune contexture of PDA following MAP Kinase inhibition is substantially missing. Inhibition of MAP Kinase, by targeting MEK1/2, has proven effective in relieving immune suppression and therefore increasing efficacy of immune checkpoint inhibitors in other solid tumors, including breast cancer and melanoma (19,20). Here, we sought to evaluate the immunological changes induced by inhibition of MEK1/2 in preclinical mouse models of PDA and found that MEK inhibition causes depletion rather than reprogramming of macrophages. Given the established role of tumor-associated macrophages in mediating resistance to therapy in PDA, we tested the effect of MEKi-induced macrophages depletion on efficacy of gemcitabine chemotherapy and observed an increased antitumor activity.

## RESULTS

### MEK inhibition results in gene expression reprogramming in mouse PDA

Using available PDA transcriptome data from the ICGC and the TCGA consortia, we found that levels of gene expression signatures indicative of MAP kinase activity (Biocarta_ERK_Pathway and MAPK Signature (21)) were higher in PDA subtypes (squamous/basal-like/quasimesenchymal) characterized by abundance of macrophage-related transcripts and immunosuppressive signaling (7,13,14) (**Figure 1A and Supplementary Figure 1A-B**). The enrichment of MAPK activity signatures in aggressive molecular subtype of PDA was also confirmed in additional 14 datasets of PDA (**Supplementary Figure 1C-D**). This data suggests that biologically aggressive molecular subtypes of human PDA have increased MAP kinase activity and led us to explore the possibility that pharmacological inhibition of MAP kinase may alter the immune contexture of PDA.

**Figure 1.**
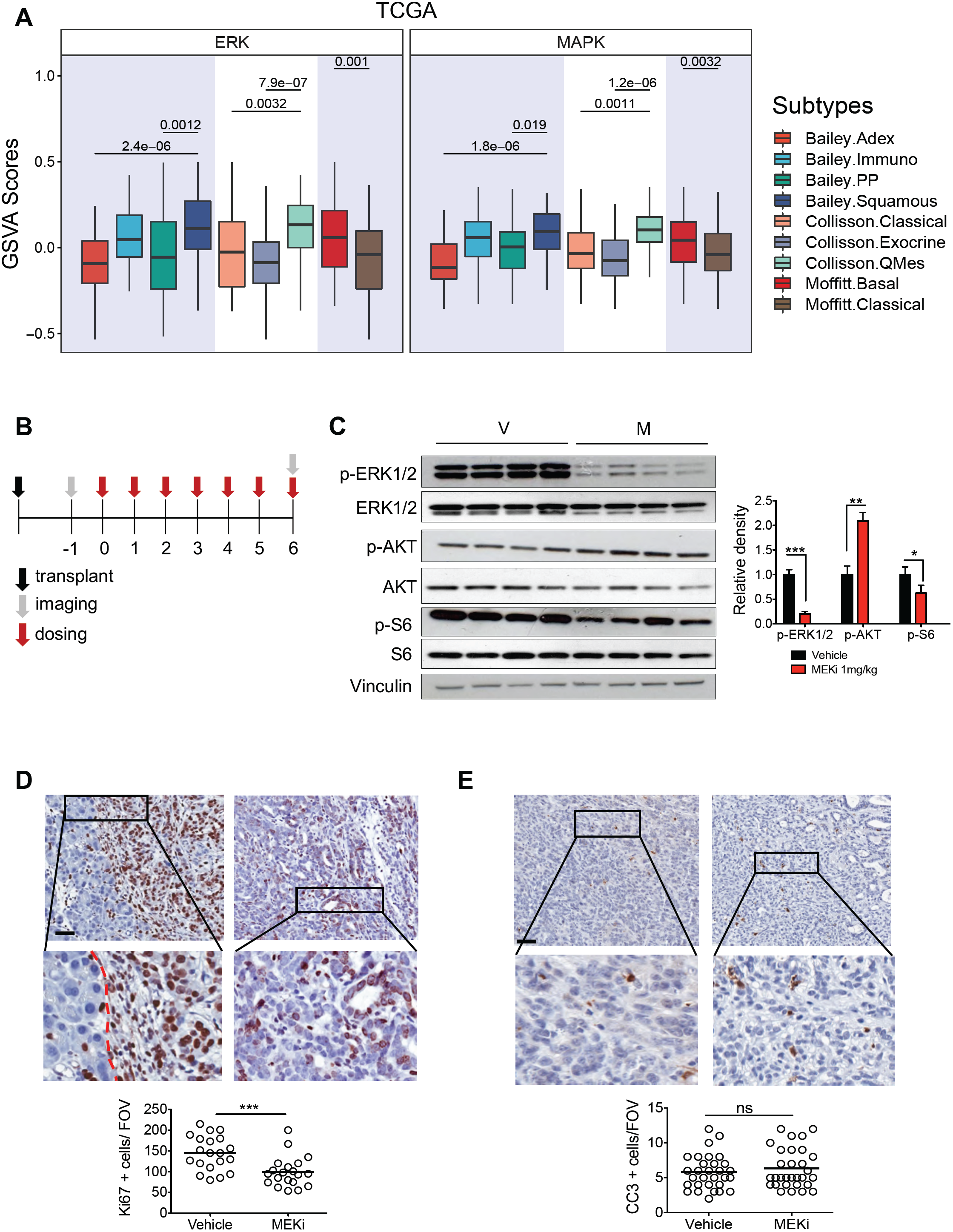
Transcriptional signatures of MAP Kinase activity in human PDA subtypes. **A** Boxplots showing the Biocarta_ERK_Pathway and the MAPK pathway (21) signatures score from GSVA stratified by Bailey (7), Moffitt (12), and Collisson (11) subtypes in the TCGA cohort. Numeric *p*-values are from Student’s *t*-test. See also Supplementary Figure 1. **B** Treatment schedule for short-term *in vivo* studies with two different doses of MEKi (0.3 and 1mg/kg) given by *oral gavage* daily. **C** Immunoblot analysis of selected signaling effectors of the MEK and AKT pathways in whole lysates from tumors treated with vehicle (V, n = 4) and 1mg/kg MEKi (M, n = 4) for 7 days. Vinculin was used as loading control. Quantification of changes in phosphorylated levels of the selected proteins after treatment is provided on the right. Error bars indicate SEMs. *, *p*<0.05; **, *p*<0.01; and ***, *p*<0.001 relative to vehicle control by Student’s *t*-test. See also Supplementary Figure 2. **D** Representative images of immunohistochemical staining for the proliferation marker Ki67 of vehicle treated (left panel) and MEKi treated tumors (right panel). Quantification is given below and was performed by counting Ki67 positive neoplastic cells in 5 field of visualization (FOV) for each mouse (n = 4/arm). ***, *p*<0.001, Student’s *t*-test. Dashed red line separates normal pancreas (left) from PDA (right). Scale represents 50 μm; insect magnification is 600x. **E** Representative images of immunohistochemical staining for the apoptotic marker cleaved caspase 3 (CC3) of vehicle treated (left panel) and MEKi treated tumors (right panel). Quantification is given below and was performed by counting CC3 positive neoplastic cells in 5 FOV for each mouse (n = 5/arm). ns, not significant Student’s *t*-test. Scale represents 50 μm; insect magnification is 600x.

First, we generated orthotopic isografts using a KPC derived cell line (FC1199) transduced with a lentiviral vector expressing luciferase for monitoring of *in vivo* growth and definition of optimal MEK1/2 inhibitor (trametinib, MEKi) dosage. Ten days following transplantation, mice were treated daily, for 7 consecutive days, with vehicle and two different concentrations of trametinib (0.3 mg/kg and 1mg/kg) (**Figure 1B**). While both dosages of MEKi delayed tumor growth (**Figure S2A**), only the higher dose of MEKi (1mg/kg) was able to effectively inhibit pathway’s flux as determined by the significant reduction in the phosphorylation of ERK1/2 (**Figure 1C and Supplementary Figure 2B**). In keeping with this, compensatory hyperphosphorylation of AKT was only observed in mice treated with 1mg/kg of MEKi (**Figure 1C**). Despite the effective inhibition of the target, the effects of MEKi on tumors were primarily cytostatic as shown by the reduction in the fraction of proliferating neoplastic cells (Ki67 positive cells, *p*<0.001) in MEKi-treated compared to vehicle-treated tumors (**Figure 1D**). Consistently, there was no clear induction of apoptosis in MEKi-treated tumors as assessed by immunohistochemical staining of cleaved caspase 3 (CC3) (**Figure 1E**).

A second array of isografts (n=17) was then generated from non-transduced KPC cell lines to monitor the effect of MEK inhibition on the immune contexture of mouse PDA by a multidimensional approach (**Figure 2A**). Following orthotopic transplantation of FC1199 cells into syngeneic mice, tumor growth was monitored by high-contrast ultrasonography (**Figure S2C**). In line with previous observations, the treatment with MEKi delayed tumor growth both after 1 and 2 weeks of treatment (**Figures S2D-E**). To evaluate in an unbiased manner the effect of MEKi on transcriptional programs of mouse PDA, we performed RNA-seq analysis and evaluated differentially expressed genes in vehicle-treated (n=4) compared with MEKi treated (n=4) FC1199 tumors at 7 days post-treatment (**Figure S2F**).

**Figure 2.**
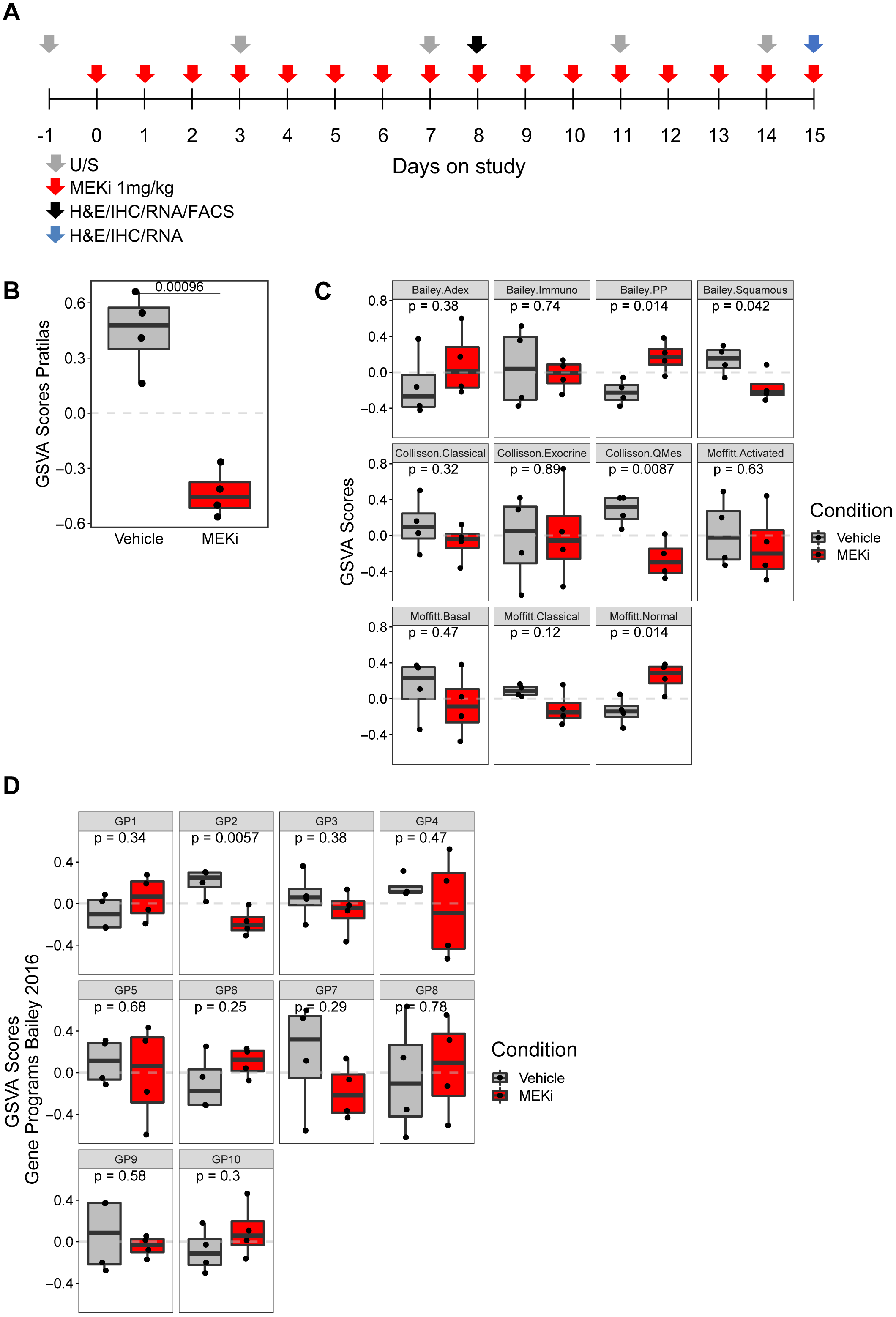
MEK inhibition results in gene expression reprogramming of mouse PDA. **A** Treatment schedule for *in vivo* studies with MEKi (trametinib, 1mg/Kg) given by *oral gavage* daily for 14 consecutive days. U/S, high-contrast ultrasonography; H&E, hematoxylin & eosin staining; IHC, immunohistochemistry; RNA, mRNA analysis either by RNA-seq or Nanostring assay; FACS, cytofluorimetric analysis. **B** Boxplot of the MAPK GSVA scores in tumors according to the indicated treatment. Numeric *p*-values are from Student’s *t*-test. **C** Boxplots of signature scores defined by Bailey (7), Moffitt (12), and Collisson (11) in tumors according to the indicated treatment. Numeric *p*-values are from Student’s *t*-test. Immuno, immunogenic; PP, pancreatic progenitor; QMes, quasimesenchymal. **D** Boxplot showing the relative enrichment of gene programs (GP) in tumors according to the indicated treatment. Numeric *p*-values are from Student’s *t*-test.

We found a total of 257 genes differentially expressed between MEKi and vehicle (fold change ≥1.5, adjusted *p*<0.05, **Supplementary Table 4**). As expected, expression of MAP Kinase associated genes was significantly reduced in MEKi treated mice (**Figure 2B**). Since we found that the levels of two independent MAP Kinase activity mRNA signatures are differentially enriched in PDA subtypes, we addressed whether MEKi induced changes in gene programs associated to human PDA molecular subtypes. FC1199-tumors were classified as squamous/basal-like/quasimesenchymal (**Figure 2C**), and one week of MEKi treatment induced a switch away from those towards the pancreatic progenitor subtype (**Figure 2C**). Moreover, inhibition of MAP kinase was also associated to increased expression of genes of the normal stromal subtype as defined by Moffitt (12) (**Figure 2C**). When looking at specific gene programs (7), there was a significant decrease in the expression of squamous gene programs (GP2, **Figure 2D**).

### MEK inhibition induces changes in the immune landscape of murine PDA

Pharmacological and genetic depletion of myeloid cells (i.e., macrophages or neutrophils) in autochthonous model of PDA has been shown to dramatically alter gene programs that specify molecular subtypes (13,14). Investigation of immune-related cell signatures in bulk RNA sequencing has proven useful to estimate the composition of immune cell populations as well as transcriptional programs underpinning mechanisms of immune regulation (7,22). Therefore, we exploited RNA-seq data and additionally performed Nanostring analysis of 700 immune-related genes on tissues from tumor-bearing mice treated for 7 and 14 consecutive days with MEKi (**Figures S3A-C**). Through interrogation of gene signatures indicative of different myeloid cell types (7,23,24) (**Supplementary Table 2**), we found a statistically significant reduction in the level of signatures driven by tumor associated macrophages (TAM) (23) **(Figure 3A and 3B**) both at 1 and 2 weeks of MEKi, while one week of MEKi was associated with elevated expression of genes related to PMN (polymorphonuclear leukocytes) (**Figure 3A and 3B**). Consistently, genes related to macrophage recruitment (*Ccl2, Cxcl3*), macrophage differentiation (*Csf3, Cxcl12*) or macrophage immunosuppressive activity (*Mrc1, Ccl17* and *Ccl22*) were among the most downregulated genes in MEKi treated tumors either at one week or two weeks of treatment (**Figure 3C and 3D**). Genes encoding for the chemokine Cxcl5 and its receptor Cxcr2 were instead amongst the most upregulated genes following one week of MEKi (**Figure 3C and 3D**). Details about shared and unique genes up- or down-regulated comparing expression profiles from mice treated for one or two weeks with MEKi is provided in **Figure S3B-E**, and **Supplementary Table 6**.

**Figure 3.**
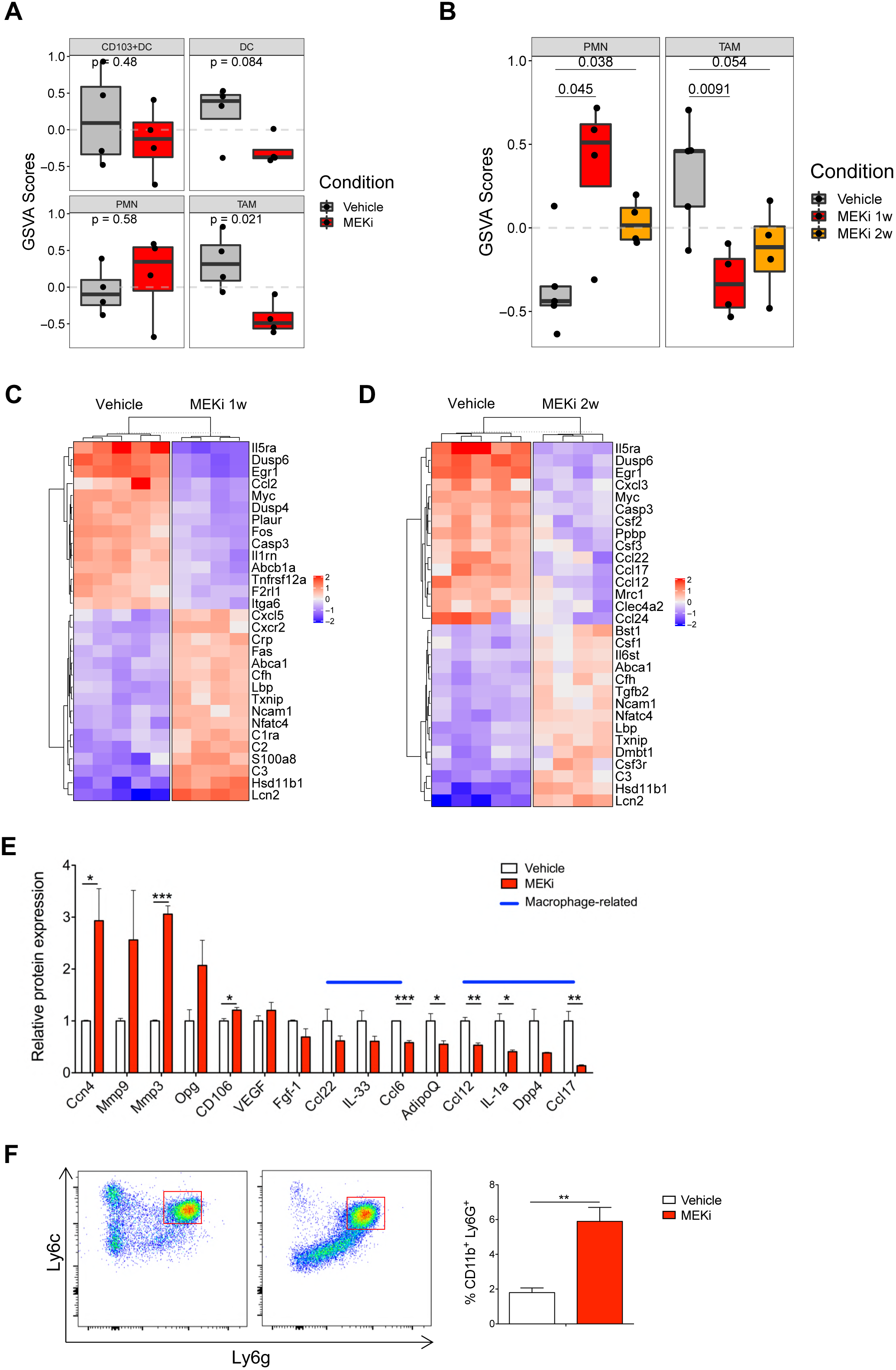
MEKi treatment of murine PDA isografts is associated with gene expression changes indicative of altered infiltration by myeloid cells. **A** Boxplots of immune-related signatures (RNA-Seq data) in tumors according to the indicated treatment. **B** Boxplots of immune-related signature scores (Nanostring data) in tumors according to the indicated treatment. Numeric *p*-values are from Student’s *t*-test. Please refer to Supplementary Table 2 for details on genes used to define the specific gene signatures. TAM, tumor-associated macrophages; PMN, polymorphonuclear neutrophils; DC, dendritic cells. **C** Heatmap showing changes in the expression pattern of 30 most differentially expressed immune-related genes in the comparison between: vehicle and one-week of MEKi. **D** Heatmap showing changes in the expression pattern of 30 most differentially expressed immune-related genes in the comparison between: vehicle and two-weeks of MEKi. **E** Quantification of changes in the levels of cytokines/chemokines assayed in whole cell lysates from tumors treated with vehicle (n = 3) or MEKi (n = 3) for 7 consecutive days. **F** After gating on CD45+ and CD11b+ cells, single cell suspensions were evaluated for expression of Ly6c and Ly6g. This evaluation was performed on 4 vehicle-treated and 4 MEKi-treated mice. Data are displayed as average ± SEMs. **, *p*<0.01; by Student’s *t*-test. See also Supplementary Table 5 and Supplementary Figure S3.

In line with gene expression data, one week of MEKi reduced the levels of chemokines/cytokines (Ccl22, Il-33, Ccl6, Ccl12, Il-1α, Ccl17), which are also produced by TAM, in protein lysates from whole tumor tissues (**Figure 3E and Figure S3F**). On the contrary, MEKi increased levels of the matrix metalloproteases Mmp-3 and Mmp-9, the latter being mainly produced by neutrophils (**Figure 3E and Figure S3F**). Multiparametric FACS analysis of tumors treated for one week with MEKi confirmed a significant expansion of CD11b^+^Ly6G^high^Ly6C^neg^ cells (neutrophils) (**Figure 3F**).

To validate the MEKi-induced changes in the infiltration of tumor by myeloid cells at tissue level, we undertook immunophenotypic analysis of mice treated with MEKi. By staining both vehicle- and MEKi-treated tissues for the macrophage marker F4/80 and Maf, a marker of anti-inflammatory macrophages (25-27), we found a reduction of total macrophages after one week of MEKi treatment (**Figure 4A**) and a more dramatic reduction of Maf^+^ cells both after one and two weeks of treatment (**Figure 4B**). Immunostaining for Ly6G confirmed the increased infiltration of neutrophils after one week of MEKi, while no further infiltration was observed at 2 weeks of treatment (**Figure 4C**).

**Figure 4.**
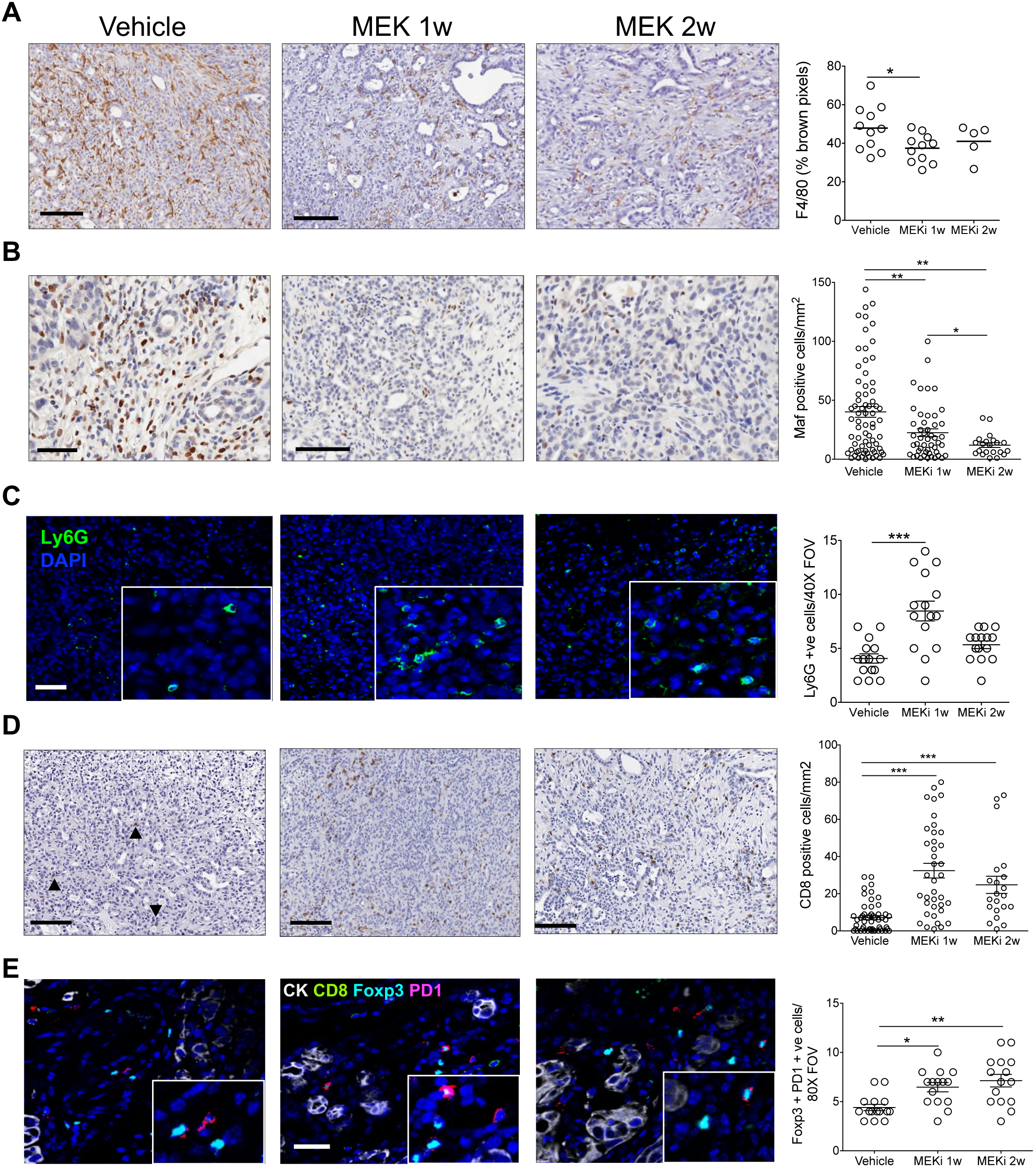
MEKi treatment of mouse PDA isografts is associated with reduction of TAM and increased infiltration of Ly6G^+^ cells. **A** Representative immunohistochemical staining for the pan-macrophage marker F4/80 in PDA tissues from mice treated with vehicle (left), one week of MEKi (middle) or two weeks of MEKi (right). Scale bars, 100μm. Quantification is provided on the right as percentage of brown pixels. **B** Representative immunohistochemical staining for the marker Maf in PDA tissues from mice treated with vehicle (left), one week of MEKi (middle) or two weeks of MEKi (right). Scale bars, 100μm. Quantification is provided on the right as number of Maf positive nuclei per mm^2^. At least 5 individual areas per case and a minimum of 4 mice/arm were evaluated**. C** Representative staining for the PMN marker Ly6G (green) in PDA tissues from mice treated with vehicle (left), one week of MEKi (middle) or two weeks of MEKi (right). Scale bar, 20 μm. Quantification is provided on the right. FOV, field of visualization. A minimum of 3 areas per case and a minimum of 3 cases were examined **D** Representative immunohistochemical staining for the T cells marker CD8 in PDA tissues from mice treated with vehicle (left), one week of MEKi (middle) or two weeks of MEKi (right). Scale bars, 100μm. Quantification is provided on the right as number of CD8 positive T cells per mm^2^. At least 5 individual areas per case and a minimum of 4 mice/arm were evaluated. **E** Multicolor immunofluorescence for pan-cytokeratin (CK, white), CD8 (green), Foxp3 (turquoise), and PD1 (magenta) in PDA tissues from mice treated with vehicle (left), one week of MEKi (middle) or two weeks of MEKi (right). Scale bar, 20 μm. Magnification is provided in insets. Nuclei were counterstained with DAPI (blue). Quantification is provided on the right. A minimum of 3 areas per case and a minimum of 3 cases were examined. *, *p*<0.05; **, *p*<0.01; and ***, *p*<0.001 by Student’s *t*-test.

In addition to rare infiltration by T cells, PDA is also characterized by poor cell priming (28-30). Therefore, we also used Nanostring data to explore transcriptional programs underpinning mechanisms of T cell function and antigen presentation. There were no significant changes induced by MEKi in immune signatures related to antigen presentation and activation of T cells (**Figure S4A**). Since appropriate evaluation of signatures related to rare cell populations might be difficult from bulk tissue expression analysis, we evaluated intratumoral infiltration of T cells populations (CD8^+^ T cells and Foxp3^+^ cells) by immunophenotyping. We found increased infiltration by CD8^+^ T cells following both 1 and 2 weeks of continuous MEKi treatment (**Figure 4D**). Multiplex immunofluorescence revealed that while the total number of Foxp3^+^ cells (Treg) were not affected (**Figure 4E and Supplementary Figure 4B**), the treatment significantly increased the fraction of Treg expressing PD1 (**Figure 4E**). PD1^+^ CD8^+^ T cells were instead rare in the tumor microenvironment of treated mice (**Figure 4E and Supplementary 4C**).

We also tested whether MEKi was also able to affect the immunogenicity of FC1199 tumors by examining the expression of MHC-I, MHC-II and PD-L1 on surface of FC119 tumor cells treated *in vitro* with MEKi for 72 hours. MEKi synergized with IFN-γ in inducing expression of MHC-I, MHC-II, and PD-L1 on the cell surface of FC1199 cells (**Figures S4D-F**), and accordingly, the combination was also able to significantly induce mRNA expression of *B2m* (**Figure S4G**). However, MEKi alone did not induce any change on the above antigen presentation markers.

### MEK inhibition induces macrophage depletion in endogenous PDA models

Next, we confirmed the MEKi-induced macrophage depletion in PDA tumor tissues in independent isografts generated with the KPC derived cell line FC1245 and in CKP mice, a highly aggressive spontaneous model of PDA (31). The reduction in the number of Maf^+^ macrophages was evident after two days and persisted after two weeks of continuous MEKi treatment of FC1245 tumor bearing mice (**Figures S5A-D).** On the contrary, two days of MEKi were sufficient to promote infiltration by neutrophils (**Figure S5E**).

To investigate effects of MEKi on macrophages on spontaneous PDA models, tumor-bearing CKP mice were treated for two days with 1mg/kg of MEKi. Consistent with the isografts models, interrogation of the differential expression of 700 immune-related genes between untreated and treated mice revealed that MEKi reduced expression of macrophage-related genes (*Ccl6, Ccl17*, and *Ccl2*) (**Figure 5A**) and of an anti-inflammatory macrophages’ signature (**Figure 5B**). Immunophenotypic analysis of CKP confirmed a significant reduction of macrophages (F4/80+ cells) and, in particular, of both CD206+ and Maf+ cells following short-term MEK inhibition (**Figure 5C**). This data suggests that MEKi has a direct effect on viability of macrophages in PDA tissues, which prompted us to investigate the sensitivity of macrophages to MEKi *in vitro*. First, we differentiated macrophages from bone-marrow cells of B6J mice and induced polarization towards both anti-tumoral (pro-inflammatory, M1-like) and pro-tumoral (anti-inflammatory, M2-like) phenotypes using combination of LPS/IFN-γ and IL-4/IL-13, respectively (**Figure S6A**). As expected, M2-like polarized macrophages displayed increased expression of anti-inflammatory markers CD206 and CD301 (**Figure S6B)** and showed preferential utilization of mitochondrial respiration compared to both M0 and M1 macrophages as previously described for M2 and tumor educated macrophages (**Figures S6C-E**) (32). M1 macrophages, on the other hand, showed increased expression of iNos and reduced mitochondrial respiratory activity compared to both M0 and M2 macrophages (**Figures S6C-E**). Next, we evaluated the sensitivity of different macrophage phenotypes to MEKi *in vitro* and found that both 2 and 3 days of MEKi treatment significantly augmented apoptosis of M2-like macrophages (**Figure 5D**). Reprogramming rather than depletion of macrophages has been recently suggested as a successful strategy to increase sensitivity towards immunoncology (33,34). Therefore, we also sought to assess if MEKi was able to induce reprogramming of macrophages towards anti-tumor (pro-inflammatory) phenotypes. To this aim, we treated polarized macrophages with subIC50 concentrations of MEKi and evaluated expression of M1-like and M2-like markers by flow cytometry to find that the treatment does not induce macrophages’ reprogramming (**Figure 6A**). In keeping with this, treatment of M2-polarized macrophages with MEKi did not revert completely their metabolic features (**Figure 6B-D**) and when global gene expression of our treated isografts was examined we did not observe increase of M1-driven signatures (**Figure S6F**).

**Figure 5.**
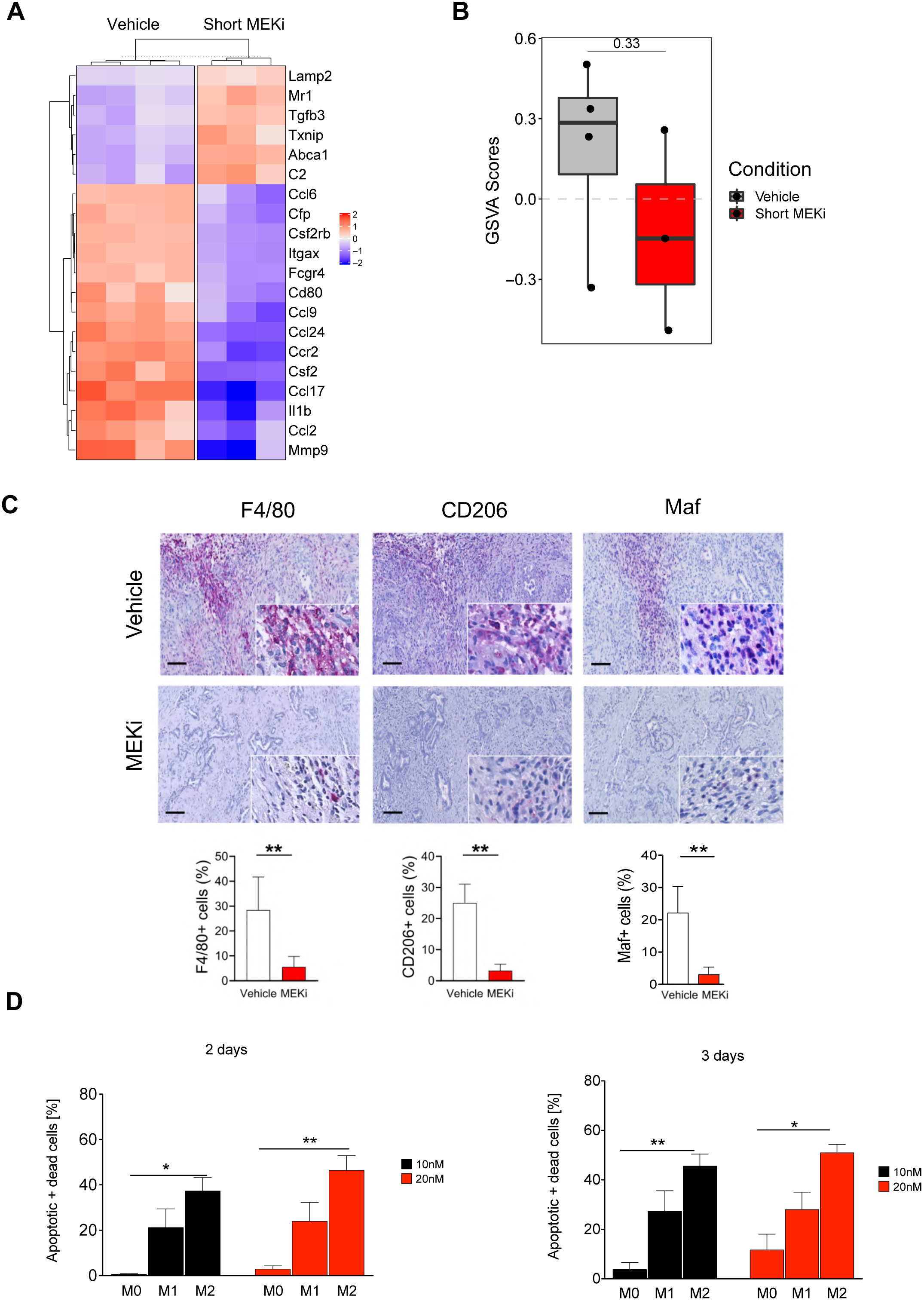
Depletion of macrophages by MEK inhibition in a spontaneous model of endogenous PDA. **A** Heatmap showing changes in the expression pattern of 30 most differentially expressed immune-related genes in the comparison between: vehicle (control) and two-days of MEKi (short MEKi). **B** Boxplots of TAM score in tumors according to the indicated treatment. Numeric *p*-values are from Student’s *t*-test. **C** Representative immunohistochemical staining for the pan-macrophage marker F4/80 (left panel), the anti-inflammatory macrophages markers CD206 and Maf (right panels) in PDA tissues from CKP mice treated for 2 days with trametinib. Quantification is provided below as percentage of positive cells relative to total cells counted in 5 high-power field for each mouse (n = 6/arm). **D** Differentially polarized macrophages were treated with MEKi (10 and 20nM) or DMSO as control for 2 and 3 days. Quantification of apoptotic cells, as percentage of apoptotic cells over untreated controls, following 2 (left panel) and 3 days (right panel) of treatment as average of 3 independent experiments. Error bars indicate standard error of the mean. **, *p*<0.01; *, *p*<0.05 by Student’s *t*-test.

**Figure 6.**
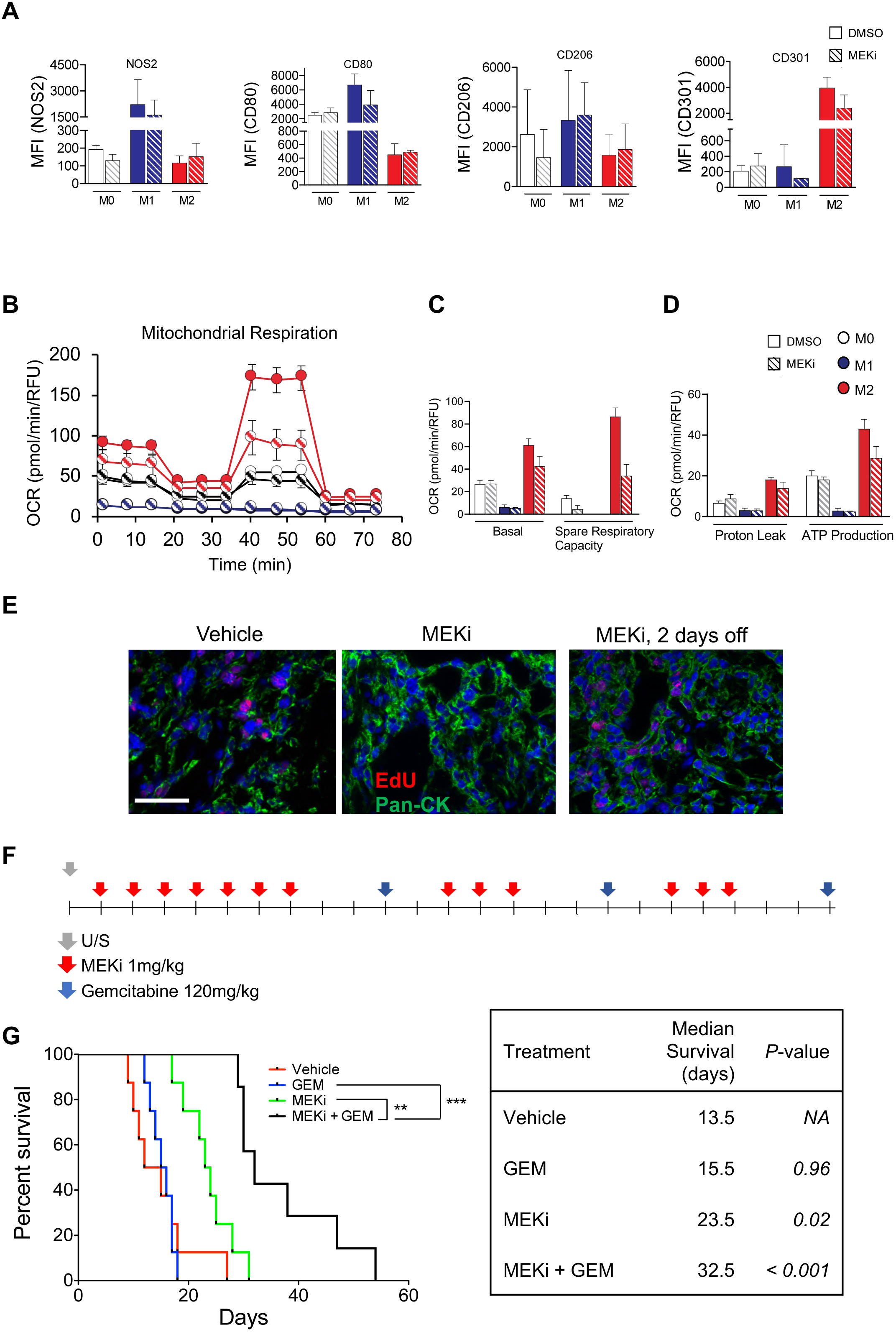
MEKi induced depletion of macrophages increases antitumor activity of gemcitabine. **A** Protein expression analysis of of M1-(Nos2, CD80) and M2-markers (CD206, CD301) of vehicle-(DMSO) and MEKi-treated macrophages. M.F.I., mean fluorescence intensity. **B** Mitochondrial respiration was detected in M0, M1 and M2 macrophages, which underwent MEKi or DMSO treatment for 30h prior to the metabolic measurement, respectively. Firstly, OCR was determined under basal conditions followed by addition of oligomycin (O, 2 µM), the uncoupler FCCP (F, 1.5 µM) and the electron transport inhibitor Rotenone/Antimycin A (R, 0,5 µM) (n = 2). The rates of basal respiration (**C**), spare respiratory capacity (**C**), proton leak (**D)**, and ATP production (**D**) were quantified. Cells were fixed and stained with DAPI and values were normalized according to respective cell numbers. **E** Representative fluorescence images of tissue staining for pan-cytokeratin (Pan-CK, green), EdU (red) and counterstained with Dapi (blue). Mice were given EdU by intraperitoneal injection 30 minutes prior to sacrifice. Scale bar, 20μm. **F** Experimental designed for survival study of tumor-bearing mice treated with vehicle, MEKi, gemcitabine (GEM), and combination of MEKi and GEM. Ultrasound was used to monitor tumor growth. **G** Survival curves of tumor-bearing mice treated as indicated in E (7-8 animals/arm). Median survival of treated mice is indicated in the table (right); *p*-value, Long-rank (Mantel-Cox) test.

Overall, we showed that MEKi treatment induces changes of the immune contexture in different murine PDA models with an early and persistent depletion of anti-inflammatory macrophages.

### MEKi-induced depletion of TAM increases response to Gemcitabine

Tumor-associated macrophages have been consistently reported to impair gemcitabine efficacy in PDA by mechanisms involving drug uptake and metabolism (15,16). Lyssiotis’s group has recently demonstrated that TAM and M2 macrophages, but not M1, interfere with gemcitabine uptake by cancer cells through release of pyrimidines (35), which is inherently linked to increased usage of mitochondrial respiration by TAM and M2 as compared to M1 macrophages.

We thus questioned how MEKi treatment might interact with standard-of-care gemcitabine-based chemotherapy given its ability of depleting the tumor microenvironment from TAM. Gemcitabine is an anticancer nucleoside that works mainly through inhibition of DNA synthesis (36). Notably, MEKi *in vivo* significantly reduced the number of cancer cells in the cell cycle (**Figure 1E**), suggesting that inhibition of MEK might reduce the number of cells targetable by gemcitabine. To determine the optimal combination schedule, we evaluated the effect of MEKi on S-phase progression following continuous drug treatment as well as after drug suspension *in vivo*. 48 hours of continuous MEKi was sufficient to block EdU uptake, while S-phase reentry of cancer cells was observed following 48 hours of MEKi discontinuation (**Figure 6E**). This result argued for alternate scheduling than concomitant treatment. Thus, we designed a survival study based on one week of continuous MEKi treatment, in order to reduce the number of M2 macrophages, followed by two days of recovery before gemcitabine administration (**Figure 6F**). While gemcitabine (Gem) monotherapy had no effect on survival of tumor-bearing mice, MEKi single agent treatment extended survival (**Figure 6G**). We observed a statistically significant increase in overall survival for tumor-bearing mice receiving gemcitabine following 7 days on, 2 days off MEKi treatment compared to MEKi (*p*=0.014) and gemcitabine (*p*=0.001) as single agents.

## DISCUSSION

Therapeutic progress beyond more intensive chemotherapeutic regimen remains an enigma in PDA with targeted and immune-based approaches failing so far. Recent sequencing efforts have uncovered actionable segments of the PDA genome (37) and revealed the existence of distinct phenotypes based on transcriptomic profiling of primary tumors (7,11,12). While showing distinct transcriptional signatures and different clinical behaviors (7,11,12), PDA subtypes all share the oncogenic activation of *KRAS* (7). Preclinical models have undoubtedly shown that oncogenic KRAS drives the generation of a complex tumor microenvironment (TME), where immunosuppressive myeloid cells represent the prominent cell type and T cells are rare (29,38). This is consistent with the definition of PDA as an immunologically “cold” tumor (30). An intense desmoplastic reaction with abundant macrophages and scant T cells is also observed in humans (39). Recent work showed that a subset of PDA do exhibit moderate infiltration of T cells (7,40) and that T cell high tumors are usually associated with reduced myeloid cells content (29).

Here, we show that transcriptional signatures of downstream Ras signaling (MAP kinase) are particularly enriched in the squamous/basal-like/quasimesenchymal subtype, which is characterized by abundance of macrophage-related transcripts, immunosuppressive signaling, and carries the worst prognosis among the human PDA subtypes. Our data are consistent with previous observations of high *KRAS* dosage and MAP kinase activity in poorly differentiated and aggressive PDA (41). As observed in several preclinical and clinical trials (42-44), effective pharmacological inhibition of MAP kinase in orthotopic isografts of KPC derived cell lines had mostly cytostatic effects. Based on mRNA profiling, our isografts models aligned with the squamous/basal-like/quasimesenchymal subtype, similarly to KPC, and therefore represents a valid experimental system for functional interrogation of the most aggressive and immunosuppressed PDA subtype. Accordingly, our model displayed extensive infiltration by macrophages and scant T cells. Integrative mRNA expression and immunophenotypic analyses showed that MEKi alters the immune contexture of PDA microenvironment and, in particular, dramatically reduces the number of tumor-associated macrophages (TAM), while promoting infiltration by neutrophils. The same MEKi-induced macrophage depletion was observed in PDA tissues from an additional KPC cell line isografts model as well as in a spontaneous aggressive model of endogenous PDA.

Moreover, we show that depletion of macrophages is seen early during the course of the *in vivo* treatment, likely due to increased drug sensitivity of M2 macrophages compared to M0 and M1. Tumor-associated macrophages are prominent mediators of T cell exclusion (14) (45). Accordingly, we found that MEKi-induced TAM depletion was associated to increased infiltration of CD8+ T cells. However, there was no indication of increased T cell reactivity or enhanced antigen-presentation by DC cells from the mRNA analysis, further supporting the hypothesis of an intrinsic DC/APC dysfunction in PDA tumor microenvironment (28). In keeping with this, *in vitro* treatment of KPC derived cell lines with MEKi failed to increase expression of genes/proteins involved in antigen presentation. A hallmark of the PDA squamous subtype is gene programs associated with squamous differentiation and macrophage function (7). A previous study has shown that depletion of macrophages in KPC using a CSF1R inhibitor resulted in “subtype switch” and reduced immunosuppressive microenvironment (14). In our model, we also observed a “switch away” from the squamous/basal-like/quasimesenchymal subtype following MEKi treatment. Taken together, our data shows that MEKi dramatically influences the TME in PDA by causing early and persistent depletion of TAM. As tumor-associated macrophages have been consistently reported to impair chemotherapy efficacy in PDA (15,16), we tested whether MEKi-induced macrophages depletion could sensitize to chemotherapy. TAMs have been shown to specifically interfere with gemcitabine uptake by cancer cells through release of pyrimidines; therefore, we used a sequence specific treatment with MEKi in combination with gemcitabine and found superior antitumor activity compared to both gemcitabine and MEKi as single agents.

Our study demonstrates that MEK inhibition induces a profound remodeling of the TME in preclinical models of PDA, which is associated to molecular subtype “switching”. Notably, we show that MEK inhibition reduces tumor infiltration by macrophages in both isografts and spontaneous mouse model of the disease, which could be exploited therapeutically to improve activity of gemcitabine.

## Acknowledgments

VC is supported by Associazione Italiana Ricerca sul Cancro (AIRC; Grant No. 18178). AS is supported by Associazione Italiana Ricerca sul Cancro (AIRC; Grant No. 12182). PD is supported by Fondazione Nadia Valsecchi (FNV) Onlus. Claudia Fiorini is supported by Fondazione Umberto Veronesi (FUV). A.S. and J.T.S. are supported by the European Union’s Seventh Framework Programme for research, technological development and demonstration (FP7/CAM-PaC) under grant agreement no. 602783. J.T.S. is also supported by the German Cancer Consortium (DKTK), and the Deutsche Forschungsgemeinschaft (DFG; KFO337/SI 1549/3-1). The funding agencies had no role in the collection, analysis and interpretation of data or in the writing of the manuscript. We thank Claudia Parolini, Paola Piccoli, and Valerio Crestan for technical assistance with immunohistochemical staining of mouse tissues. We also thank Martina De Toni for technical assistance with *in vitro* work.

## Authors’ Contribution

### Concept and design

PD, CN, DF, PFC, JTS, and VC

### Acquisition of data

DF, EF, and SDa generated isografts and performed in vivo treatment; FB performed IVIS imaging analysis; SU and FDe performed cytofluorimetric analysis; DF, SDa, and FL performed immunohistochemistry; CF and DF performed immunoblotting; MS, DF, and PFC performed mRNA analysis; RTL provided facilities.

### Analysis and interpretation of data

SB and PD performed computational and statistical analysis; MF and BR performed histological revision and assisted evaluation of immunohistochemistry; VB, DF, PD, CN, AS, PFC, JTS and VC interpreted results of experiments.

### Writing, review, and/or revision of the manuscript

DF, CN, PD, PFC, JTS, and VC co-wrote the manuscript. All authors commented on the manuscript and approved final version.

### Study supervision

DF, CN, PD, PFC, JTS, and VC

## MATERIALS AND METHODS

### Cell Cultures

The murine PDA cell lines FC1199 and FC1245 were kindly provided by the Tuveson laboratory (Cold Spring Harbor Laboratory, NY, USA) and were generated from PDA tumors of KPC (<u>K</u>ras^G12D/+^; <u>p</u>53^R172H/+^; Pdx1-<u>C</u>re) mice (46). FC1199^luc/Gfp^ cells were generated using procedures reported elsewhere (47). All cells were maintained in DMEM (Invitrogen) supplemented with 10% FBS (Gibco). Cells were routinely tested for the presence of *Mycoplasma* contamination using MycoAlert Mycoplasma Detection Kit (Lonza).

### Mice

SIx- to 8-weeks old C57Bl/6J (B6J) mice were purchased from Charles River Laboratory. All animal experiments regarding transplanted mice were conducted in accordance with procedures approved by CIRSAL at University of Verona (approved project 655/2017-PR). *Ptf1a*^*wt/Cre*^;*Kras*^*wt/LSL-G12D*^;p53^*fl/fl*^ (CKP) (31) was used as tumor model of spontaneous PDA for phenotypic characterization and quantification of immune cells following MEKi treatment. All animal protocols related to CKP mice were performed according with appropriate guidelines and the experimental protocols were approved by the local Animal Use and Care Committee at the University Hospital of Essen, Germany.

### Drug treatments

For *in vitro* experiments, trametinib (Selleck) was dissolved in DMSO, whose final concentration was less than 0.1% (v/v). For *in vivo* experiments, trametinib was dissolved in a solution of 0.5% hydroxypropyl-methylcellulose, 0.2% tween 80 and ddH_2_O (pH 8) for oral administration. Gemcitabine (Gem) was obtained from the Pharmacy of the University and Hospital Trust of Verona as pharmaceutical grade suspension at 38mg/mL, which was diluted to 12mg/mL in PBS before intraperitoneal injection (i.p.) at 120mg/kg. Cell death induction of macrophages upon MEKi treatment was evaluated using Annexin V Apoptosis Detection Kit (BD Biosciences). Polarized macrophages were incubated with or without 10 nM or 20 nM trametinib or with respective volume of DMSO (0.1 % v/v) for 48 hours and 72 hours. Macrophages were collected and washed with PBS (Thermo Fisher Scientific Inc), prior to staining with FITC Annexin V and PI (Acros Organics) according to manufacturer’s instructions. FACS analysis was conducted using BD FACSCelesta flow cytometer (BD Biosciences). Raw data were analyzed using FlowJo software version 10.5.3 (LLC).

### Orthotopic tumor models and GEMM

6- to 8-weeks old B6J mice were transplanted with FC1199 or FC1245 cells (2.5×10^5^ – 5×10^5^) resuspended in a 50% v/v solution of Matrigel in PBS. A volume of 50μL of solution was injected into the pancreatic tail. For the definition of optimal trametinib dosage, FC1199^luc/Gfp^ (5×10^5^ cells/mouse) were orthotopically transplanted in B6J mice and, 10 days post transplantation, mice were injected intraperitoneally with 200μL of a concentrated luciferin solution (15mg/mL, PerkinElmer, Inc.) for IVIS imaging analysis (PerkinElmer). Mice were treated daily by oral gavage for one week using two different concentrations of trametinib (0.3mg/kg and 1mg/kg). IVIS was performed at day −1 (the day before the beginning of treatments), then at day 6 (i.e., the day before necropsy). For therapeutic intervention studies, FC1199 (3.5×10^5^ cells/mouse) were orthotopically transplanted in B6J mice. Upon detection of a palpable mass, mice were subjected to high-contrast ultrasound imaging using the Vevo 2100 System with a MS250, 13–24 MHz scanhead (Visual Sonics, Inc, Amsterdam, NL) and enrolled in treatment studies according to the outlined treatment schedules. Mice were administered vehicle or 1mg/kg MEKi every day by oral gavage unless otherwise indicated. Gemcitabine was administered i.p. at a dose of 120mg/kg once a week. CKP mice were enrolled in treatment studies at age of 6 weeks as previously described (31). Mice were treated with vehicle or trametinib (1mg/kg) daily, by oral gavage, according to the reported treatment schedules.

### Statistical Analysis and Data mining

For data mining and pancreatic cancer subtypes stratifications we used different gene expression datasets. The two main datasets were the PACA-AU cohort of the ICGC consortium and the TCGA-PAAD cohort of the TCGA consortium. The first was downloaded from the supplementary material of the corresponding publication (7). This dataset contains normalized expression values (TMM normalized using edgeR Bioconductor package, converted to CPM and log2 transformed) of 96 pancreatic cancer patients; for subtypes stratification, Z-scores were calculated for each gene. Associated clinical data were downloaded from https://dcc.icgc.org/releases/current/Projects/PACA-AU. The second dataset represents the TCGA-PAAD cohort, downloaded from http://firebrowse.org/?cohort=PAAD, which consists of the RNA-Seq gene expression profile of 178 pancreatic cancer patients. According to other publications that disputed the purity of some samples, we restricted the samples’ number to 148. Fourteen additional microarray and RNA-Seq datasets were downloaded and used for subtype stratification. These datasets are listed in **Supplementary Table 1**. Normalized expression values were obtained with the GEOquery Bioconductor R package. For harmonization, all the datasets were Z-scores standardized before subsequent analysis. The grouping of the samples in Bailey’s, Collisson’s, and Moffitt’s subtypes (7,11,12) was performed with the GSVA Bioconductor package. The gene sets used for the stratification were retrieved from the original publications. Unless indicated, all the *p*-values refers to Student’s *t*-test.

Other conventional techniques are described in Supplementary Materials and Methods.

## FIGURE LEGENDS

**Supplementary Figure 1.**
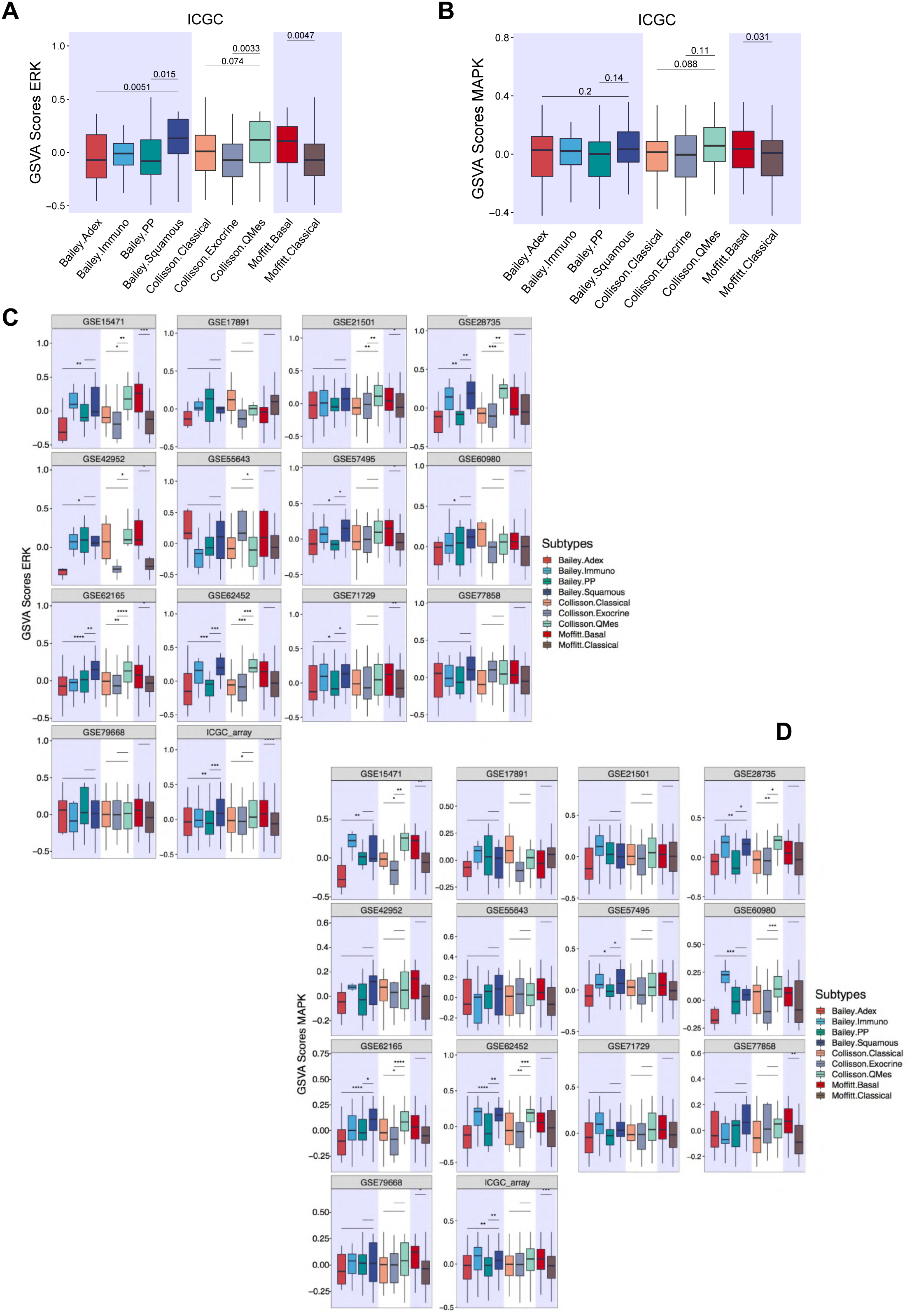
Transcriptional signatures of MAP Kinase activity in human PDA subtypes. Boxplots showing the the Biocarta_ERK_Pathway (**A**) and the MAPK pathway (21) (**B**) signatures score stratified by Bailey (7), Moffitt (12), and Collisson (11) subtypes in the ICGC cohort. Numeric *p*-values are from Student’s *t*-test. Boxplots showing the Biocarta_ERK_Pathway (**C**) and the MAPK pathway (**D**) (21) signature scores from GSVA stratified by Bailey (7), Moffitt (12), and Collisson (11) subtypes in 14 different gene expression datasets, both microarray and RNA-Seq. ****, *p*<<0.0001; ***, *p*<0.001; **, *p*<0.01, and *, *p*<0.05 as determined by Student’s *t*-test.

**Supplementary Figure 2.**
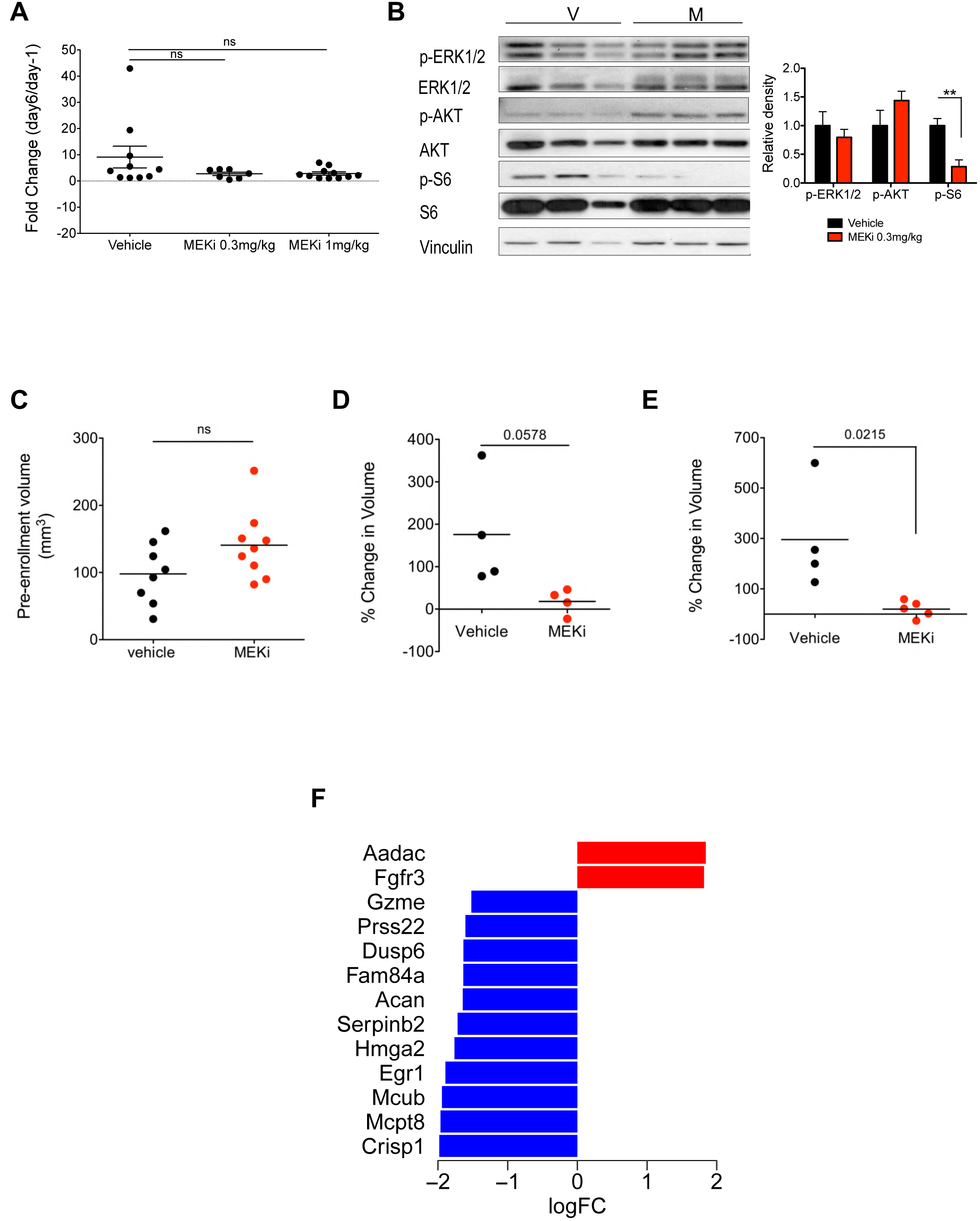
Effect of MEKi on mouse PDA isografts. **A** Scatter dot plot showing changes in bioluminescence intensities for individual mouse tumors treated with vehicle (n = 10), 0.3mg/kg of MEKi (n = 7), and 1mg/kg of MEKi (n= 11). The results are expressed as average with error bars representing SEMs. ns, not significant by Student’s *t*-test. **B** Immunoblot analysis of selected signaling effectors of the MEK and AKT pathways in lysates from mouse tumors treated with vehicle (V, n= 3) and 0.3mg/kg MEKi (M, n = 3) for 7 days. Vinculin was used as loading control. Quantification of changes in phosphorylated levels of the selected proteins after treatment is provided on the right. Error bars indicate SEMs. **, *p*<0.01 relative to vehicle control by Student’s *t*-test. **C** Scatter dot plot showing pre-enrollment tumor volumes as measured by ultrasound indicating no significant differences between initial tumor volumes in the two cohorts of tumor-bearing mice. **D** Scatter dot plot showing changes in the volumes of individual mice (expressed as percentage of the initial tumor volume) after one week of treatment in the two cohorts. *p*-values, Student’s *t*-test. **E** Scatter dot plot showing changes in the volumes of individual mice (expressed as percentage of the initial tumor volume) after two weeks of treatment in the two cohorts. *p*-values, Student’s *t*-test. **F** Differential expression analysis of the transcriptomic response to 7 days of MEKi treatment (shown are the 13 most significantly modulated genes with logFC ≥ 1.5).

**Supplementary Figure 3.**
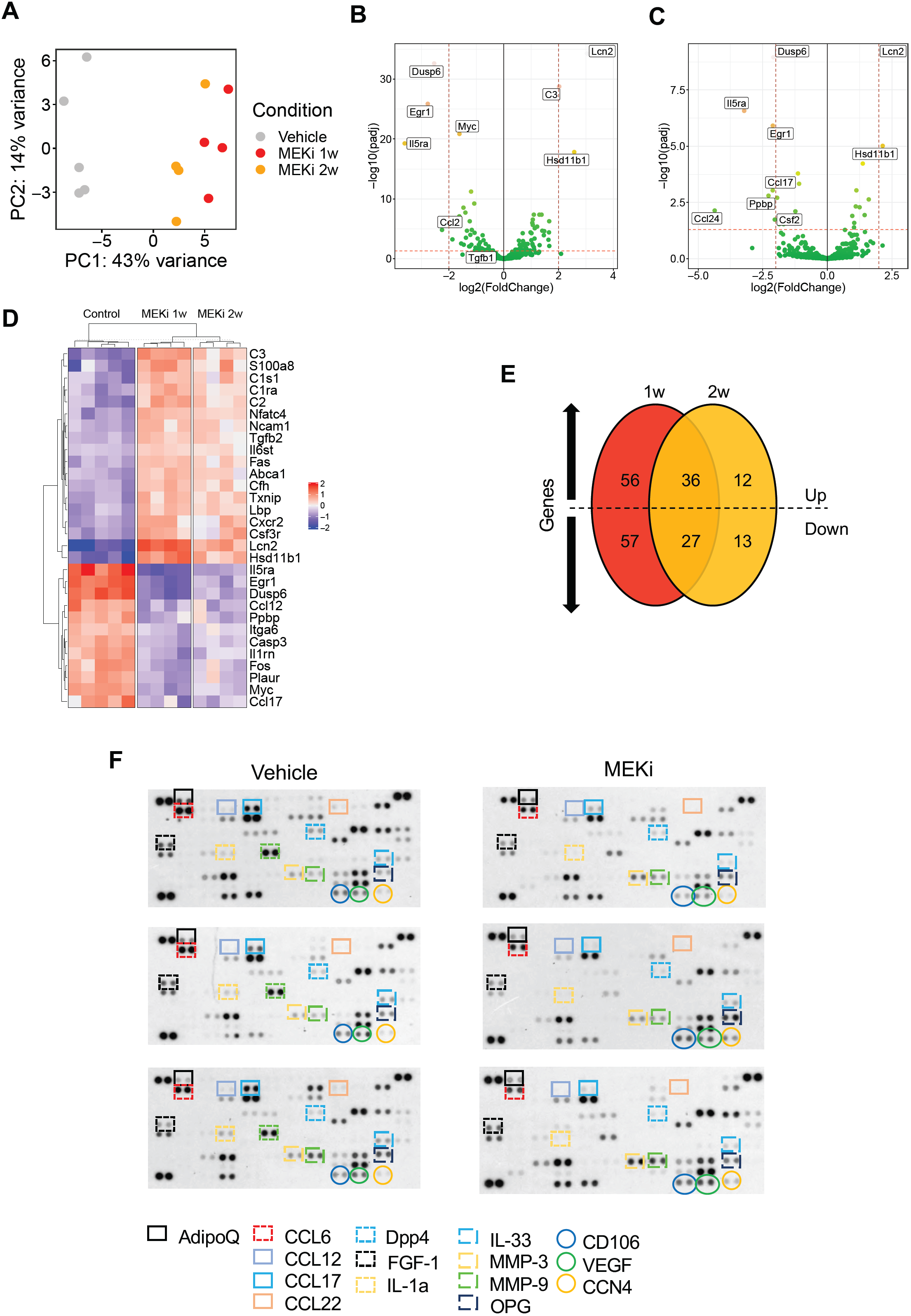
MEKi elicits changes in the transcriptome and proteome of murine PDA. **A** Principal component analysis of gene expression data for tumors treated with vehicle, 7 days of MEKi, and 14 days of MEKi. **B** Volcano plot of differences in gene expression (NanoString platform) between vehicle- (n = 5) and one-week MEKi-treated tumors (n = 4). Indicated are the genes with log_2_FC expression ≥ 2 and adjusted *p*<0.05. **C** Volcano plot of differences in gene expression (NanoString platform) between vehicle- (n = 5) and two-weeks MEKi-treated tumors (n = 4). Indicated are the genes with log_2_FC expression ≥ 2 and adjusted *p*<0.05. **D** Heatmap showing normalized and centered *vst* counts from DESeq2 package among tumors treated with vehicle, 7 days of MEKi, and 14 days of MEKi. **E** Venn diagram illustrates shared and unique genes upregulated and downregulated in the comparison of expression profiles of tumors treated with MEKi for one week (red) two weeks (orange) compared to vehicle treated tumors; please refer to Supplementary Table 6 for details. **F** Mouse Cytokine arrays for tumors treated with vehicle or MEKi for 7 consecutive days. Colored rectangles highlight mouse cytokine quantified in Figure 3E.

**Supplementary Figure 4.**
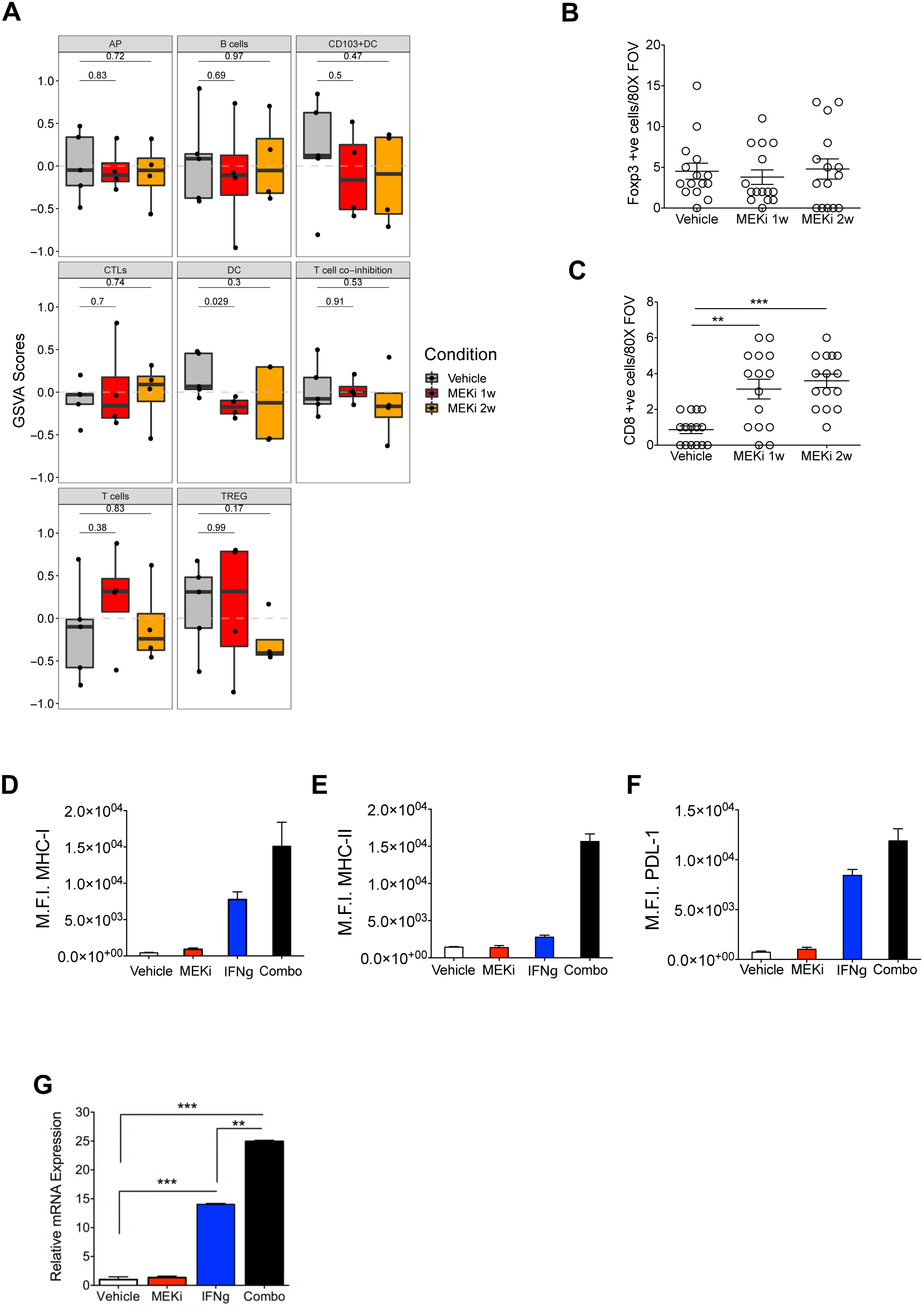
Effect of MEKi on antigen presentation. **A** Boxplots of immune-related signature scores in tumors according to the indicated treatment. Numeric *p*-values are from Student’s *t*-test. Please refer to Supplementary Table 2 for details on genes used to define the specific gene signatures. TREG, regulatory T cells; CTLs. Cytotoxic T lymphocytes; AP, antigen presentation; DC, dendritic cells. **B** Quantification of Foxp3+ cells from Figure 4E. **C** Quantification of CD8+ cells from Figure 4E. ***, *p*<0.001; and **, *p*<0.01 by Student’s *t*-test. Single cell suspension from FC1199 cells treated with vehicle, MEKi, IFN-γ and combination of MEKi and IFN-γ were evaluated for the cell surface expression of MHC I (**D**), MHC II (**E**), and PD-L1 (**F**). Data are displayed as average of two biological replicates. M.F.I., mean fluorescence intensity. **G** Changes in mRNA levels of *B2m* in FC1199 cell lines following 72 hours treatment with MEKi and IFN-γ (IFNg) as single agents or in combination. Levels of *B2m* were normalized to *Hprt1*, and then levels of gene following treatment were normalized to those of the vehicle control. Error bars indicate SEMs. ***, *p*<0.001; and **, *p*<0.01 by Student’s *t*-test.

**Supplementary Figure 5.**
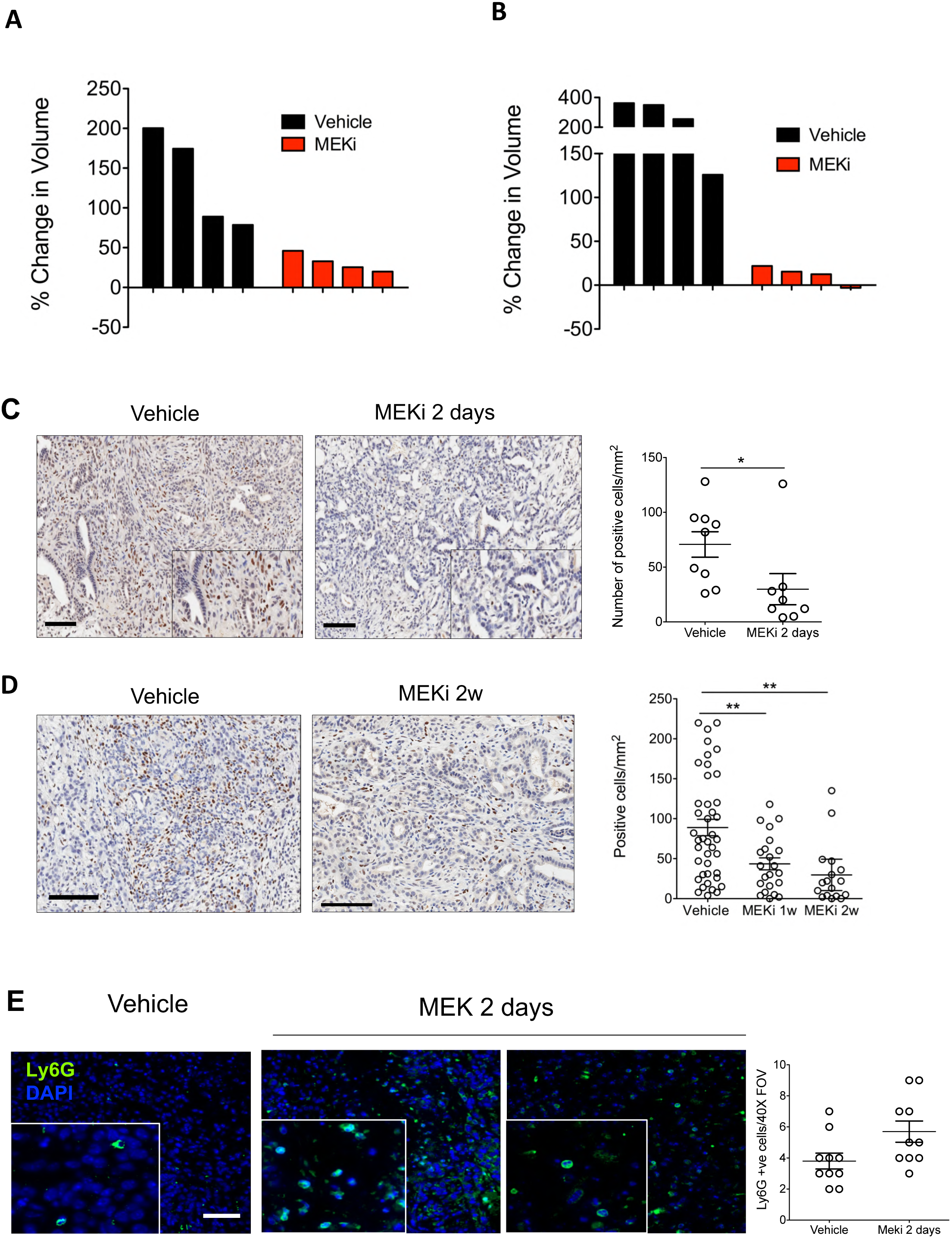
Validation of MEKi-induced TAM depletion in an additional isografts model. **A** Waterfall plot of individual tumor volume changes as measured by ultrasound after one week of treatment. **B** Waterfall plot of individual tumor volume changes as measured by ultrasound after two weeks of treatment. **C** and **D** Representative immunohistochemical staining for the marker Maf in PDA tissues from mice treated with vehicle, two days and two weeks (2w) of MEKi. Scale bars, 100μm. Quantification is provided on the right as number of Maf^+^ nuclei per mm^2^. At least 5 individual areas per case and a minimum of 4 mice/arm were evaluated for the long term-treatment, while 3 individual areas per case and 3 mice/arm were evaluated for the short-term treatment. **E** Immunofluorescence staining for Ly6G (green) in tissues from tumor-bearing mice treated with vehicle or MEKi for 2 days. Scale bar, 20 μm. Quantification is provided on the right. FOV, field of visualization. *, *p*<0.05 by Student’s *t*-test.

**Supplementary Figure 6.**
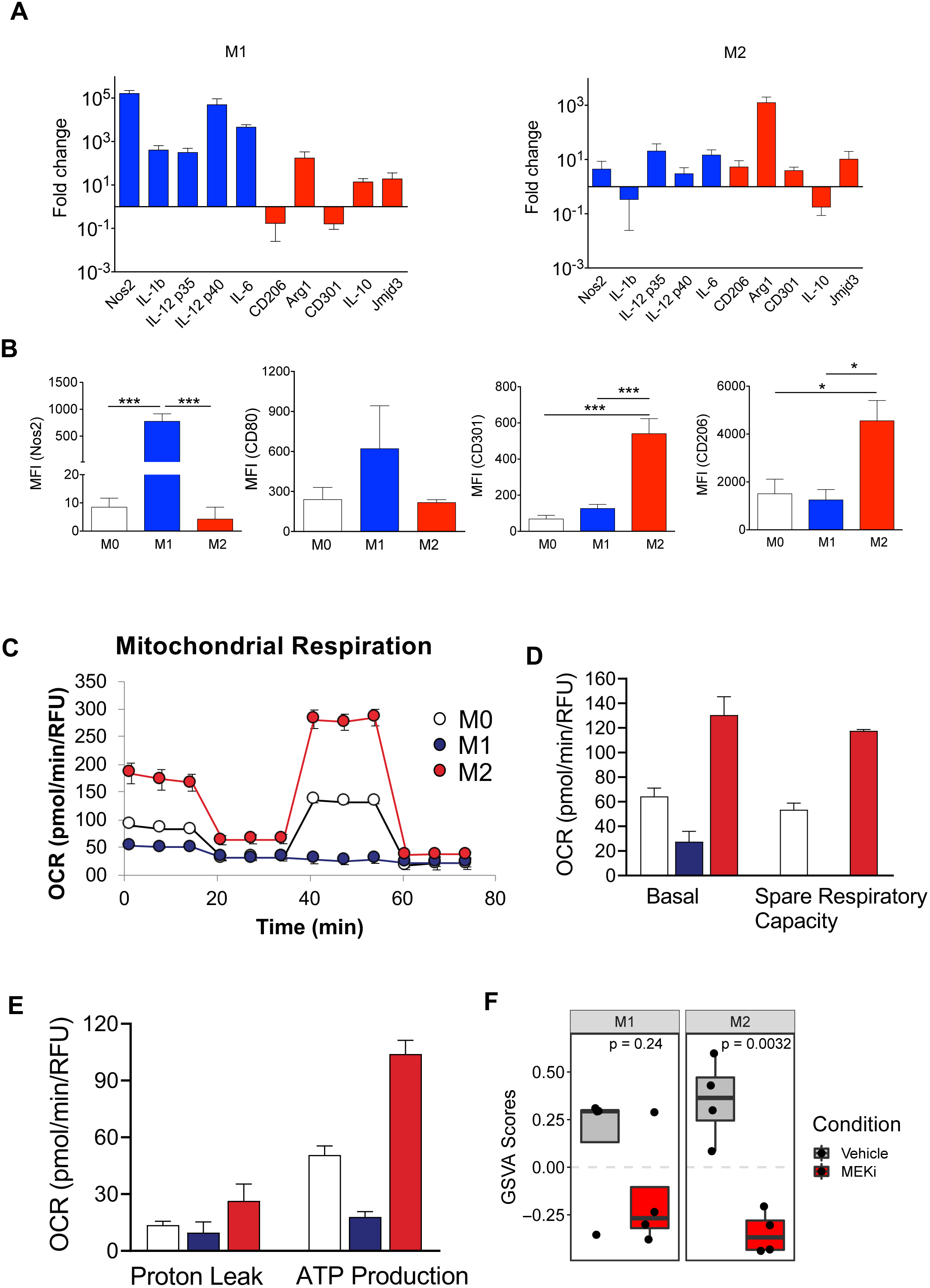
Characterization of *in vitro* generated bone marrow-derived macrophages. Bone-marrow derived progenitor cells were differentiated into unstimulated macrophages (M0) and afterwards polarized into pro-inflammatory (M1) or anti-inflammatory (M2) macrophages. **A** Gene expression analysis of multiple pro- and anti-inflammatory marker genes by qRT-PCR. Fold change expression of M1 and M2 macrophage subsets were calculated by normalizing γCT values to the unstimulated M0 state. Data are displayed as average of 3 independent experiments. Error bars indicate standard deviation. **B** Protein expression analysis of M1- (Nos2, CD80) and M2-markers (CD206, CD301) to validate successful polarization. M.F.I., mean fluorescence intensity. *, *p*<0.05, ***, *p*<0.001 by Student’s *t*-test. **C** Mitochondrial respiration reflected by OCR levels was detected in M0, M1 and M2 macrophages under basal conditions followed by addition of oligomycin (O, 2µM), the uncoupler FCCP (F, 1.5µM) and the electron transport inhibitor Rotenone/Antimycin A (R, 0,5µM) (n = 2). **D** and **E** the rates of basal respiration, spare respiratory capacity, proton leak, and ATP production were quantified. Cells were fixed and stained with DAPI and values were normalized according to respective cell numbers. **F** Boxplots of M1 and M2 signatures in tumors according to the indicated treatment. Numeric *p*-values are from Student’s *t*-test.

