## Supplementary Methods for "MAP Kinase inhibition reshapes tumor microenvironment of mouse pancreatic cancer by depleting anti-inflammatory macrophages"

**Supplementary Materials and Methods**

**Isolation of bone marrow–derived cells and macrophage differentiation**

Bone marrow–derived cells were isolated from femur and tibia of healthy B6J mice aged between 8 and 12 weeks. Bone marrow was collected by flushing bones with RPMI medium (Life Technologies). Red blood cells (RBC) were lysed with RBC Lysing Buffer (Sigma-Aldrich GmbH) for 7 minutes at room temperature. Cells were then washed and resuspended in macrophage medium (RPMI, 20% FBS, 1% NEAA, 1% sodium pyruvate; Life Technologies) containing 50 ng/mL murine M-CSF (PeproTech) for 5 days. Macrophage purity were assessed based on CD11b and F4/80 expression using flow cytometry. To induce M1-like polarization, cells were treated with 1 μg/mL LPS (Sigma-Aldrich) and 40 ng/mL of IFNγ (PeproTech) for 24 hours, whereas for M2-like phenotype, macrophages were treated with 40 ng/mL IL4 (PeproTech) and 40 ng/mL of IL-13 (Peprotech) for 48 hours.

**Metabolic extracellular flux analysis**

Macrophage metabolism was investigated by measuring oxygen consumption rate (OCR) using Seahorse XFe96 Analyzer (Agilent Technologies Inc.) with Seahorse XF Cell Mito Stress Test Kit (Agilent) following manufacturer’s instructions. Bone marrow-derived macrophages were seeded in XF-96 cell culture plate (Agilent Technologies Inc.) and allowed to attach overnight. Cells were treated with 2 μM Oligomycin, 1,5 μM FCCP as well as 0,5 μM Rotenone/Antimycin A and subsequent changes in OCR were used to calculate OXPHOS characteristics. For investigation of inhibitor-induced subset-specific metabolic changes, macrophages were seeded overnight in XF-96 cell culture plate and treated with 10 nM Trametinib or respective volume of DMSO for 30 hours and then analyzed immediately by Seahorse XF Cell Mito Stress Test Kit (Agilent) following manufacturer’s instructions. Following OCR measurements, cells were fixed in 4 % PFA for 10 min at RT and stained with DAPI for normalization according to respective cell numbers. DAPI intensity was measured using Tecan (Tecan Trading AG). Normalization and data analysis were performed using Seahorse Wave Desktop 2.6 software.

**Histology and Immunohistochemistry**

Tissues were fixed in 10% neutral buffered formalin and embedded in paraffin. Sections were subjected to Hematoxylin and Eosin as well as immunohistochemical staining. The following primary antibodies were used for immunohistochemical staining of PDA isografts: F4/80 (ab6640, Abcam); MAF (sc-7866, Santa Cruz); CD8 (14-0808-82, Thermo Scientific), Cleaved Caspase-3 (9661, Cell Signaling), and Ki67 (D2H10, Cell Signaling). Slides were scanned at 20x magnification and digitalized using the Aperio Scan-Scope XT Slide Scanner (Aperio Technologies). Quantification of F4/80 staining was performed using ImageJ.

For multiplex immunofluorescence staining, we used the Opal Multiplex IHC Kit (Akoya) and the following antibodies: pan-keratin (4279S, Cell Signaling) 1:100, CD8α (98941, Cell Signaling) 1:100; FoxP3 (12653, Cell Signaling) 1:100; and PD1 (84651S, Cell Signaling) 1:100. Briefly, FFPE sections were deparaffinized and then subjected to several sequential of microwave treatment and staining. Each includes antigen retrieval by heat-induced epitope retrieval using citrate buffer (pH6), a protein blocking followed by primary antibody, introduction of Secondary-HRP, and incubation with Opal Fluorophore for 10 min at room temperature. After all sequential staining reactions, sections were counterstained with DAPI (Vector lab). Slides were scanned by Leica TCS SP5 laser scanning confocal (Leica) with 80× objective magnification and digitalized by the Leica Application Suite X (LAS X) software. Immunofluorescence images were quantified using at least 3 fields per tumour. Immunostaining of CKP derived tissues was performed using Dako REAL^TM^ Alkaline Phosphatase Detection System (Dako, Germany), following manufacturer’s instructions. Antigen retrieval of formalin-fixed paraffin-embedded (FFPE) sections was achieved by heat-induced epitope retrieval using citrate buffer (pH6) for CD11b (ab133357, Abcam), CD206 (ab64693, Abcam), and MAF (sc-7866, Santa Cruz) and Tris/EDTA (pH9) for CD163 (ab182422, clone EPR19518, Abcam), and proteinase K treatment for F4/80 (T-2028, clone BM8, BMA biomedicals). Following blocking with serum free protein solution (Dako) for 1h at RT, slides were incubated with respective primary antibody for 30 min at RT, and then subjected to FastRed chromogen development. Slides were scanned and digitalised using Zeiss Axio Scanner Z.1 (Carl Zeiss AG, Germany) with 2,5x and 10x magnification. 5 representative images from each tissue section of each mouse were used to determine the percentage of positive cells from IHC staining using Definiens software (Definiens AG, Germany). For quantification, total cells number of every tissue section was determined based on nuclei staining detected by the software. Areas of extensive immune cell infiltration, e.g. infiltrating lymph nodes or tertiary lymphoid structures, were avoided.

**Immunoblot**

Protein lysates were obtained from fresh-frozen tissues by dissolving up to 25 14µm cryostat sections in RIPA lysis buffer. Tissue lysates were separated in 4%–12% Bis-Tris NuPage gels (Life Technologies). Western blots were probed with the following antibodies: Akt (9272), Phospho-Akt (Ser473) (4060), p44/42 MAPK (Erk1/2) (9102), phospho-p44/42 MAPK (Erk1/2) (Thr202/Tyr204) (4376), Phospho-S6 Ribosomal protein (Ser 235/236) (4858), S6 Ribosomal protein (54D2). Vinculin (4650) was used as loading control. All antibodies were from Cell Signaling. The immunoblots were visualized with ECL plus (Amersham/GE Healthcare Europe GmgH, Munchen, Germany).

**Flow cytometry**

Flow cytometry-based immune phenotype of tumors was performed according to already published protocols (1). Briefly, tumors were minced and digested for about 1 hour at 37 °C under continuous mixing with a digestive mix containing 1mg/mL collagenase IV, 0.1mg/mL hyaluronidase, and 30U/mL DNAse in RPMI 1640, all purchased from Sigma-Aldrich. The cell suspension was separated from the undigested material using a 70-μm cell strainer (Corning). One million of cells were incubated with anti-mouse CD16/32 (Biolegend) and subsequently stained with the following antibodies according to the vendor’s instructions: CD11b (M1/70, Thermo Fisher Scientific), CD11c (N418, Thermo Fisher Scientific), LY6C (HK1.4, Biolegend), LY6G (1A8, Biolegend), CD45 (30F11, Biolegend), For analysis of cell lines, single cell suspension were probed with fluorochrome conjugated mAb anti-mouse MHC I (28-8-6, Biolegend), MHC II (AF6-120.1, EBioscience), CD274 (10F.9G2, Biolegend). Samples were acquired on a FACS Canto II (BD Biosciences, USA) and analyzed with FlowJo software (FlowJo LLC, USA). Multi-color flow cytometry of differentiated macrophages tissues was performed using BDFACSCelesta (BD Biosciences) using the following antibodies: iNOS (clone CXNFT, eBioscience); CD80 (clone 16-10A1, BioLegend); CD206 (clone C068C2, BioLegend); CD301 (clone LOM-14, BioLegend).

**Cell cycle analysis**

To determine the effect of MEKi on cell-cycle progression *in vivo*, 3 groups of 5 tumour-bearing mice were treated with 1mg/kg of trametinib or vehicle by oral gavage daily for 2 days. The mice were sacrificed, and tumours removed at 2 and 48 hours after the final dose. Tumor-bearing mice were given 50mg/kg of the thymidine analogue EdU (Invitrogen) by intraperitoneal injection. Percentage of neoplastic cells in S-phase was evaluated by immunofluorescence staining of frozen tumour section using the Click-iT EdU Alexa Fluor 647 Imaging Kit (Invitrogen).

**Cytokine array**

Protein Profiler Mouse XL Cytokine Array (ARY028, R&D) was used. Tumors (treated as outlined) were frozen and protein lysates obtained according to the manufacturer’s instructions; 300µg of lysates were incubated with array over night at 4 °C. Multiple exposures were used to determine optimal sensitivity and films were scanned using trans-illumination. Data was quantified using ImageJ.

**qRT-PCR analysis**

RNA was extracted with Trizol®Reagent (Life Technologies) method. 1µg of DNase-treated RNA was reverse transcribed using SuperScript® VILO^TM^ cDNA Synthesis Kit (Life Technologies) in a volume of 20µl according to the manufactures instructions. Samples were diluted to a final concentration of 10ng/µl. TaqMan was performed in duplicate or triplicate using 20ng of cDNA and the following TaqMan® probe (TaqMan® Gene Expression Assay): *B2m* (Mm00437762_m1). *Hprt1* was used as reference gene. Relative gene expression quantification was performed using the ΔΔCt method with the Sequence Detection Systems Software, Version 1.9.1 (Applied Biosystems). Expression of genes related to different macrophage phenotypes was conducted at University Hospital of Essen and performed using the Roche LightCycler 480 with the SYBR green I Master Kit (Roche GmbH). Primers for qRT-PCR were designed using NCBI Primer Blast and purchased from Eurofins MWG Operon GmbH. Primers’ sequences can be found in **Supplementary Table 3**. Data were analyzed using ΔCT calculations where Cyclophilin A served as reference control for normalization. Amplification efficacy was experimentally determined or assumed as 2. Fold change expression of respective macrophage subsets were calculated by normalizing ΔCT values to the unstimulated M0 state. For fold change expression after inhibitor treatment, ΔCT values of vehicle treated subsets were subtracted from ΔCT values of respective inhibitor treated subsets.

**RNA sequencing**

RNA was extracted from PDA cell cultures or freshly isolated tissues (isografts) using TRIzol (Invitrogen), followed by column-based purification with the PureLink RNA Mini Kit (Ambion). RNA from pancreas of CKP mice was isolated using the Qiagen RNAeasy isolation kit (Qiagen). The quality of purified RNA samples was determined using a Bioanalyzer 2100 (Agilent) with an RNA 6000 Nano Kit. RNAs with RNA Integrity Number (RIN) values greater than 7.5 were used to generate sequencing libraries using the TruSeq Stranded Total RNA Kit (Illumina) per manufacturer’s instructions. Sequencing libraries were run at Eurofinsgenomics using an Illumina HiSeq2000. Sequence pairs were mapped to mouse genome GRC38, using the STAR 2.5.3a software (2) and the annotations retrieved from gencode vM16. Raw counts were normalized using DESeq2 v1.18.1 rlog function (3). The first principal component of the mean-adjusted gene expression matrix was strongly associated with acini contamination. The contamination was confirmed by reviewing cryostat sections of the corresponding specimens; therefore, this component was removed using SVD factorization and recomposition. Gene differential expression analysis was performed using the limma v3.34.2 R package (4). Geneset enrichment analysis was performed using the GSVA v1.26.0 R package (5). Differential enrichment between MEKi- and vehicle-treated samples was tested for selected gene sets using Student’s *t*-test. The list of gene sets used is available in the **Supplementary Table 2a.** For analysis of macrophage populations, we used the experimentally-derived gene signatures described in Jablonski et al (6).

**Nanostring Analysis**

Expression analysis was performed using Nanostring Mouse PanCancer Immune Profiling Kit (Diatech XT-CSO-MIP1-12). 25 ng of total RNA was used according to the manufacturer’s protocol. The matrix of counts was then used for differential expression analysis with DESeq2 Bioconductor package (3). DESeq2 was used in combination with RUVSeq (7) in order to control housekeeping genes expression and for normalization purposes. A batch factor of variation was calculated from the expression of the housekeeping genes, and such factor was then added to the DESeq2 design formula. Gene set variant analysis was performed with the GSVA Bioconductor package with the following parameters: method = 'gsva', mx.diff = TRUE, kcdf = 'Gaussian' (5). The list of gene sets used, and associated references is available in the **Supplementary Table 2b**.
