## Supplementary Table 3 for "MAP Kinase inhibition reshapes tumor microenvironment of mouse pancreatic cancer by depleting anti-inflammatory macrophages"

**Supplementary Table 3. List of primers’ sequences used for qRT-PCR analysis**

| **Primer** | **Sequence** |
| --- | --- |
| qmArg1_for | ATCGGAGCGCCTTTCTCAAA |
| qmArg1_rev | GGTCTCTCACGTCATACTCTGT |
| qmIL12p35_for | TCCCGAAACCTGCTGAAGAC |
| qmIL12p35_rev | CTGGTTTGGTCCCGTGTGAT |
| qmIL12p40_for | GAGTGGGATGTGTCCTCAGAA |
| qmIL12p40_rev | GTCCAGTCCACCTCTACAACA |
| qmIL1b_for | TGCCACCTTTTGACAGTGATG |
| qmIL1b_rev | ATGTGCTGCTGCGAGATTTG |
| qmIL6_for | ACTTCACAAGTCGGAGGCTT |
| qmIL6_rev | TGCAAGTGCATCATCGTTGT |
| qmJmjd3_for | CTGTAGCCCATAGGACCCAC |
| qmJmjd3_rev | GTCTCCGCCTCAGTAACAGC |
| qmMgl1_for | GGAAGCCAAGACTTCACACTG |
| qmMgl1_rev | CTGGACGGAAACCAAGACAC |
| qmCD206_for | AACAAGAATGGTGGGCAGTC |
| qmCD206_rev | TTTGCAAAGTTGGGTTCTCC |
| qmiNOS_for | CGTGAAGAAAACCCCTTGTGC |
| qmiNOS_rev | GGAACATTCTGTGCTGTCCCA |
| qmIL10_for | AGGCGCTGTCATCGATTTCT |
| qmIL10_rev | ATGGCCTTGTAGACACCTTGG |
| qmCyclophilinA_for | ATGGTCAACCCCACCGTG |
| qmCyclophilinA_rev | TTCTGCTGTCTTTGGAACTTTGTC |
